# Copper Blocks V-ATPase Activity and SNARE Complex Formation to Inhibit Yeast Vacuole Fusion

**DOI:** 10.1101/625517

**Authors:** Gregory E. Miner, Katherine D. Sullivan, Chi Zhang, Logan R. Hurst, Matthew L. Starr, David A. Rivera-Kohr, Brandon C. Jones, Annie Guo, Rutilio A. Fratti

## Abstract

The accumulation of Copper in organisms can lead to altered functions of various pathways, and become cytotoxic through the generation of reactive oxygen species. In yeast, cytotoxic metals such as Hg^+^, Cd^2+^, and Cu^2+^ are transported into the lumen of the vacuole through various pumps. Copper ions are initially transported into the cell by the copper transporter Ctr1 at the plasma membrane and sequestered by chaperones and other factors to prevent cellular damage by free cations. Excess copper ions can subsequently be transported into the vacuole lumen by an unknown mechanism. Transport across membranes requires the reduction of Cu^2+^ to Cu^+^. Labile copper ions can interact with membranes to alter fluidity, lateral phase separation and fusion. Here we found that CuCl_2_ potently inhibited vacuole fusion by blocking SNARE pairing. This was accompanied by the inhibition of V-ATPase H^+^ pumping. Deletion of the vacuolar reductase Fre6 had no effect on the inhibition of fusion by copper. This suggests that that Cu^2+^ is responsible for the inhibition of vacuole fusion and V-ATPase function. This notion is supported by the differential effects chelators. The Cu^2+^-specific chelator TETA rescued fusion, whereas the Cu^+^-specific chelator BCS had no effect on the inhibited fusion.

---

Divalent cations play numerous roles in cellular maintenance and membrane trafficking. Among these cations, calcium is well known for its role in signaling via calmodulin and calcineurin, and regulating synaptic vesicle trafficking, while divalent metals such as zinc and copper are less understood. Copper is an essential trace metal that functions in aerobic respiration, superoxide dismutase activity and iron acquisition (1). Yet, excess free copper leads to toxicity and is associate with Wilson’s disease and liver failure (2). Many of the deleterious effects of elevated labile copper is the generation of reactive oxygen species (ROS) which can damage membranes, proteins and DNA, and exacerbate neurodegenerative diseases such Alzheimer’s and Huntingtin’s (3, 4). Aside from the generation of ROS, copper can interact with membranes to affect lateral phase separation, membrane fluidity, and can block exocytosis in PC-12 cells (5–9). With respect to lysosomes, brief copper exposure leads to the transport of the copper pump ATP7B from the plasma membrane to lysosomes and can stimulate the exocytosis of degradative enzymes (10).

In yeast, labile copper ions are transported into the vacuole lumen to reduce toxicity. Copper can subsequently be transported of copper out of the vacuole upon its reduction from Cu^2+^ to Cu^+^ by Fre6 (11). Cu^+^ is exported from the vacuole through the Ctr2 transporter, and its deletion results in the hyper-accumulation of vacuolar copper (12). Although the mechanism for copper export from the vacuole is known, the importer of the metal remains unclear. It is possible that copper import is carried out by a broad spectrum transporter and not a high affinity pump.

Due to the effects of elevated copper on membranes in mammalian cells, we hypothesized that yeast vacuole dynamics could also be affected by this cation. In this study we found that addition of exogenous Cu^2+^ strongly inhibited vacuolar fusion. An examination of the distinct stages of membrane fusion indicated that Cu^2+^ inhibited *trans*-SNARE complex formation and H^+^ uptake by the V-ATPase. The effects of copper were likely due to the Cu^2+^ form as specific chelators rescued fusion, while Cu^+^ chelators had no effect.

## Results and Discussion

### Copper inhibits vacuole fusion

Copper has been shown block the membrane fusion of chromaffin granules in PC-12 cells (6), while it stimulates lysosome exocytosis in HeLa cells (10). Here we asked if copper would alter *in vitro* yeast vacuole homotypic fusion. To start, we added a concentration curve of CuCl_2_ to vacuole fusion reactions at the start of the incubation period. We found that vacuole fusion was blocked in a dose-dependent manner with an IC_50_ of 15 μM **(Fig. 1A)**. Copper has also been shown to interact with the vacuolar TRP family Ca^2+^ channel Yvc1 (13). To test whether the inhibitory effect of Cu^2+^ on vacuole fusion was associated to this channel, we used vacuoles from *yvc1*Δ yeast. As seen in **Figure 1B**, the inhibitory effect of copper on *yvc1*Δ vacuoles was nearly indistinguishable from its effect on wild type vacuoles. The lack of a difference indicates that the effect of Cu^2+^ was independent of Yvc1. To verify whether the effect of CuCl_2_ on vacuole fusion was specific we tested other divalent cations as well as a second copper salt. As shown in **Figure 1C**, vacuole fusion was resistant to 100 µM ZnCl_2_, MgCl_2_, CoCl_2_, while vacuoles were equally sensitive to both CuCl_2_ and CuSO_4_ **(Fig. 1C).** We have also tested Ca^2+^ at 100 µM and see no effect on fusion (not shown). These data indicate the effects of copper on vacuole fusion were specific to the metal and not due to a charge effect. While Ca^2+^ can inhibit vacuole fusion, it only does so at millimolar concentrations (14). This makes the effect of copper at micromolar concentrations of particular interest.

**Figure 1.**
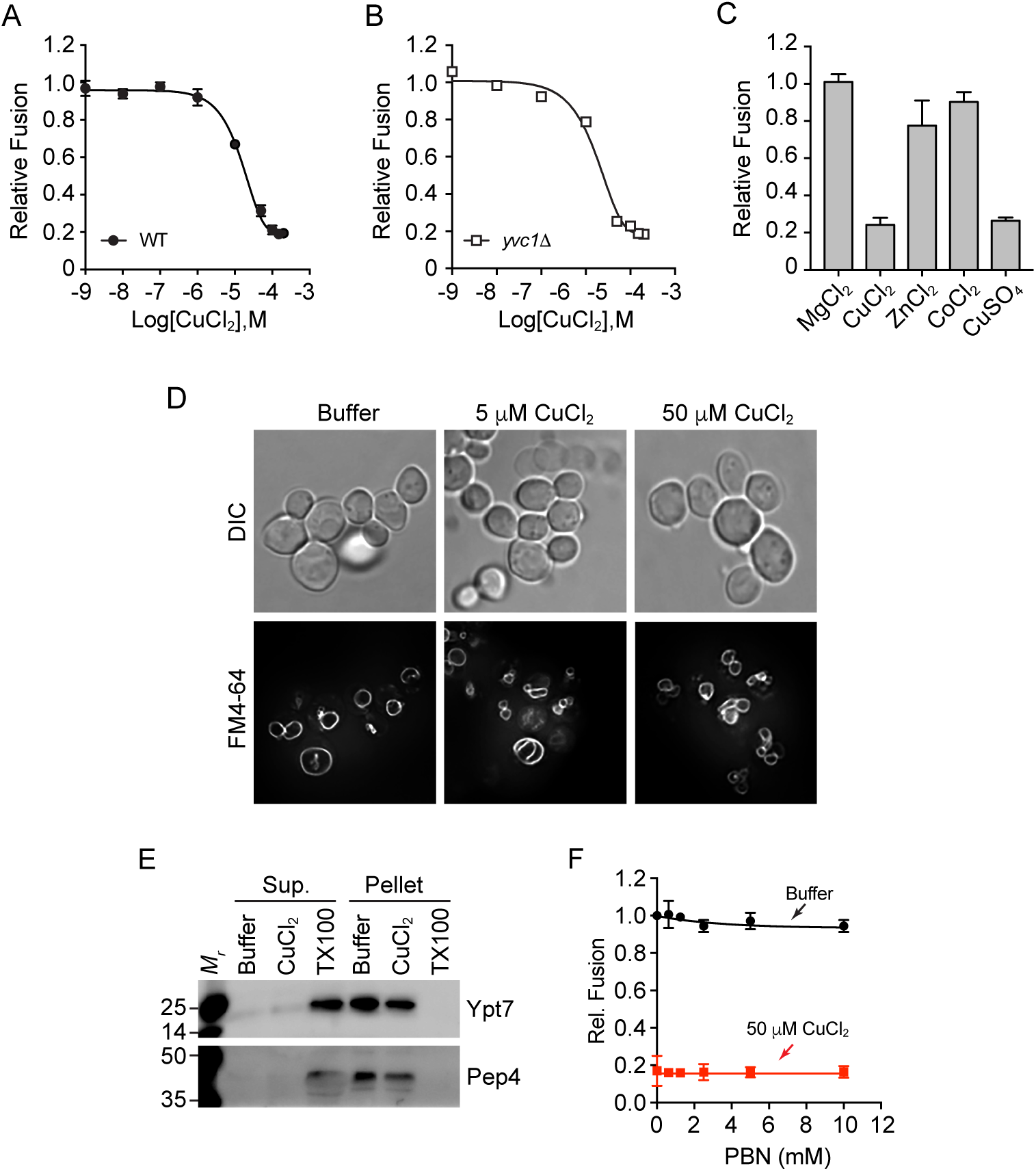
CuCl_2_ inhibits vacuole homotypic fusion. Vacuoles isolated from wild type (BJ3505 and DKY6281) **(A)** or *yvc1*Δ (RFY74-75) **(B)** fusion reporter strains were incubated with buffer alone or a concentration curve of CuCl_2_. **(C)** Wild type vacuoles were incubated with 100 µM MgCl_2_, ZnCl_2_, CoCl_2_, CuCl_2_, or CuSO_4_. Fusion reactions were incubated for 90 min at 27°C. After incubation membranes were solubilized and incubated with *p*-nitrophenyl phosphate to measure Pho8 activity. *p-*nitrophenolate was measured at OD_400_. Fusion values were normalized to the untreated control set to 1. **(D)** BJ3505 cells were grown to log phase and treated with CuCl_2_. Vacuoles were visualized by staining with FM4-64 and cells were visualized using DIC. **(E)** DKY6281 vacuoles were incubated with 100 µM CuCl_2_, 0.1% TX-100 or buffer for 90 min at 27°C. After incubation, the soluble and membrane fractions were separated by centrifugation (16,000 x g, 10 min, 4°C), mixed with SDS-loading buffer and resolved by SDS-PAGE. The soluble luminal protease Pep4 and the membrane anchored Ypt7 were probed for by immunoblotting. **(F)** Wild type vacuoles were treated with or without CuCl_2_ in the presence of a concentration curve of PBN to eliminate oxygen radicals. Error bars are S.E.M. (n=3).

The inhibition of in vitro vacuole fusion is often accompanied by in vivo vacuole fragmentation. To test the effects acute copper treatment we incubated log-phase yeast with 0, 5 or 50 µM CuCl_2_. The vacuoles were stained with the vital dye FM4-64 and vacuole morphology was examined by fluorescence microscopy and whole cells were visualized with DIC. In **Figure 1D** we show that acute CuCl_2_ treatment led to vacuole fragmentation. This is in accord with our previous results, further indicating that CuCl_2_ inhibits vacuole fusion at low micromolar levels.

### Copper does not inhibit vacuole fusion through reactive oxygen species generation

Copper ions can lead to the production of oxygen radicals that damage membranes through lipid peroxidation culminating in vesicle lysis. To test whether the effect of CuCl_2_ on vacuole fusion was due to membrane lysis we looked for the release of the soluble luminal protease Pep4. Fusion reactions were incubated with reaction buffer alone, 100 µM CuCl_2_, or 0.5% Triton X-100 (TX100). After incubation, vacuoles were pelleted through centrifugation and separated from the supernatant. Pellets were resuspended with a starting volume of reaction buffer after which both pellet and supernatant fractions were mixed with equal volumes of SDS-loading buffer. Proteins were resolved by SDS-PAGE and probed by Western blotting for Pep4 and the membrane anchored protein Rab GTPase Ypt7. We found that copper did not result in the release of Pep4 from the vacuole lumen. Rather, Pep4 remained in the pellet with Ypt7 **(Fig. 1E)**. As expected Triton X-100 solubilized both proteins that only appeared in the supernatant fraction. This indicates that copper did not cause vesicle lysis and that its effect on fusion was due to its effect on the fusion machinery itself.

To further verify that CuCl_2_ was not leading to the production of oxygen radicals, we used the free radical spin trap *N-tert*-butyl-α-phenylnitrone (PBN), which has been shown to reduce free radicals produced in the presence of copper (15). We added a concentration curve of PBN to reactions containing 50 µM CuCl_2_ or buffer alone. This showed that PBN, even at 10 mM, was unable to restore fusion in reactions treated with copper **(Fig. 1F)**. This suggests that copper does not inhibit vacuole fusion through creating oxygen radicals.

### Transport of Copper ions into the vacuole lumen

We next asked whether copper ions were transported into the vacuole lumen. One way to test this is to track resistance to an external copper chelator. The Cu^2+^ selective chelator Triethylenetetramine (TETA) was chosen for this experiment (16). TETA also chelates Zn^2+^, however, this was not relevant as Zn^2+^ had no effect on vacuole fusion. In **Figure 2A** we show that adding equimolar TETA and CuCl_2_ to vacuoles fully restored fusion, indicating that Cu^2+^ was directly responsible for the effect on fusion. Next, we added TETA at different time points after the addition of CuCl_2_ to vacuoles. TETA quickly lost its ability to block the effect of CuCl_2_ on fusion, suggesting that Cu^2+^ was no longer available to bind the chelator **(Fig. 2B)**. A likely reason would be that copper ions were transported into the vacuole lumen.

**Figure 2.**
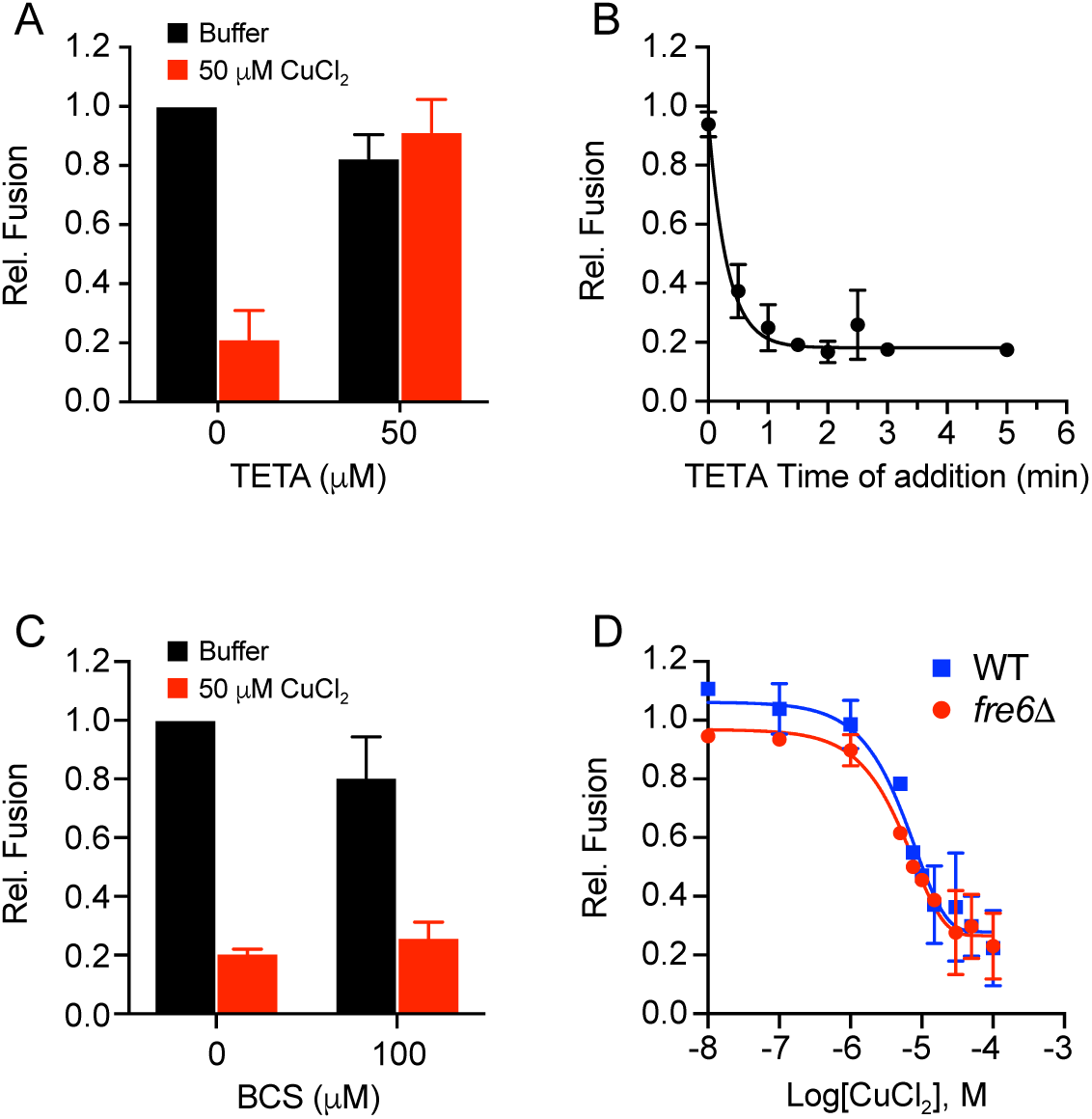
Cu^2+^ inhibits vacuole fusion. **(A)** Fusion reactions were incubated with or without CuCl_2_ and treated with TETA to chelate cupric (Cu^2+^) ions. **(B)** Fusion reactions containing 50 µM CuCl_2_ were supplemented with TETA at the indicated times. Fusion reactions were incubated for a total of 90 min at 27°C before developing. **(C)** Fusion reactions were incubated with or without CuCl_2_ and treated with BCS to chelate cuprous (Cu^+^) ions. **(D)** Wild type and *fre6*Δ vacuoles were incubated with a concentration curve of CuCl2 and tested for fusion after 90 min of incubation at 27°C. Error bars are S.E.M. (n=3).

Although much is known about the transport of copper from the vacuole lumen to the cytosolic side of the vacuole through Ctr2, the identity of a specific copper importer remains unknown. Copper transport by Ctr2 at the vacuole or Ctr1 at the plasma membrane requires that copper is reduced to Cu^+^. At the vacuole copper is reduced by Fre6 (11). Because of the relationship of cation reduction and transport, we tested whether copper was being reduced as part of its effect on fusion. To test if Cu^+^ was generated as part of its inhibition of fusion, we used the Cu^+^ chelator Bathocuproine disulfonate (BCS) (17). Two BCS molecules are needed to chelate one Cu^+^, thus 100 µM BCS was used to chelate 50 µM CuCl_2_. Unlike the effect of TETA in restoring vacuole fusion, we found the BCS was unable to rescue fusion in the presence of 50 µM CuCl_2_ (**Fig. 2C**). This suggested that copper was not being reduced outside the lumen as part of its inhibitory effect on vacuole fusion. We also examined whether Cu^2+^ reduction was necessary to inhibit fusion by deleting Fre6. In **Figure 2D**, we show that *fre6*Δ vacuole fusion was equally sensitive to CuCl_2_ as compared to the wild type parent strains. These data suggest that the effects of copper on vacuole fusion are carried out by the Cu^2+^ cation.

### Cu^2+^ inhibits vacuole fusion after the priming stage

While we know that Cu^2+^ inhibits vacuole fusion, the specific stage at which it exerts its effect is unclear. Vacuole fusion traverses several stages that begins with the activation of SNAREs through a process termed priming. This stage occurs when the AAA+ protein Sec18 is recruited to inactive *cis*-SNARE complexes that are present on each membrane and prohibited from interacting with their cognate SNAREs on other membranes. Sec18/NSF associates with *cis*-SNARE complexes through its adaptor protein Sec17/α-SNAP. Sec18 hydrolyses ATP to disrupt *cis*-SNARE bundles resulting in the release of Sec17 from the membrane (18). Thus, priming can be measured by the loss of Sec17 from the membrane. Vacuoles were incubated with reaction buffer, 100 µM Cu^2+^ or 1 mM *N*-ethylmaleimide (NEM), an inhibitor of Sec18/NSF function (19, 20). Individual reactions were incubated at increasing time increments after which they were centrifuged to separate the membrane bound (pellet) and solubilized (supernatant) fractions of Sec17. These experiments showed that Cu^2+^ did not negatively affect SNARE priming, as similar levels of Sec17 were released relative to the buffer control **(Fig. 3A-B)**. By contrast, SNARE priming was fully inhibited by NEM, as demonstrated by the complete block of Sec17 release.

**Figure 3.**
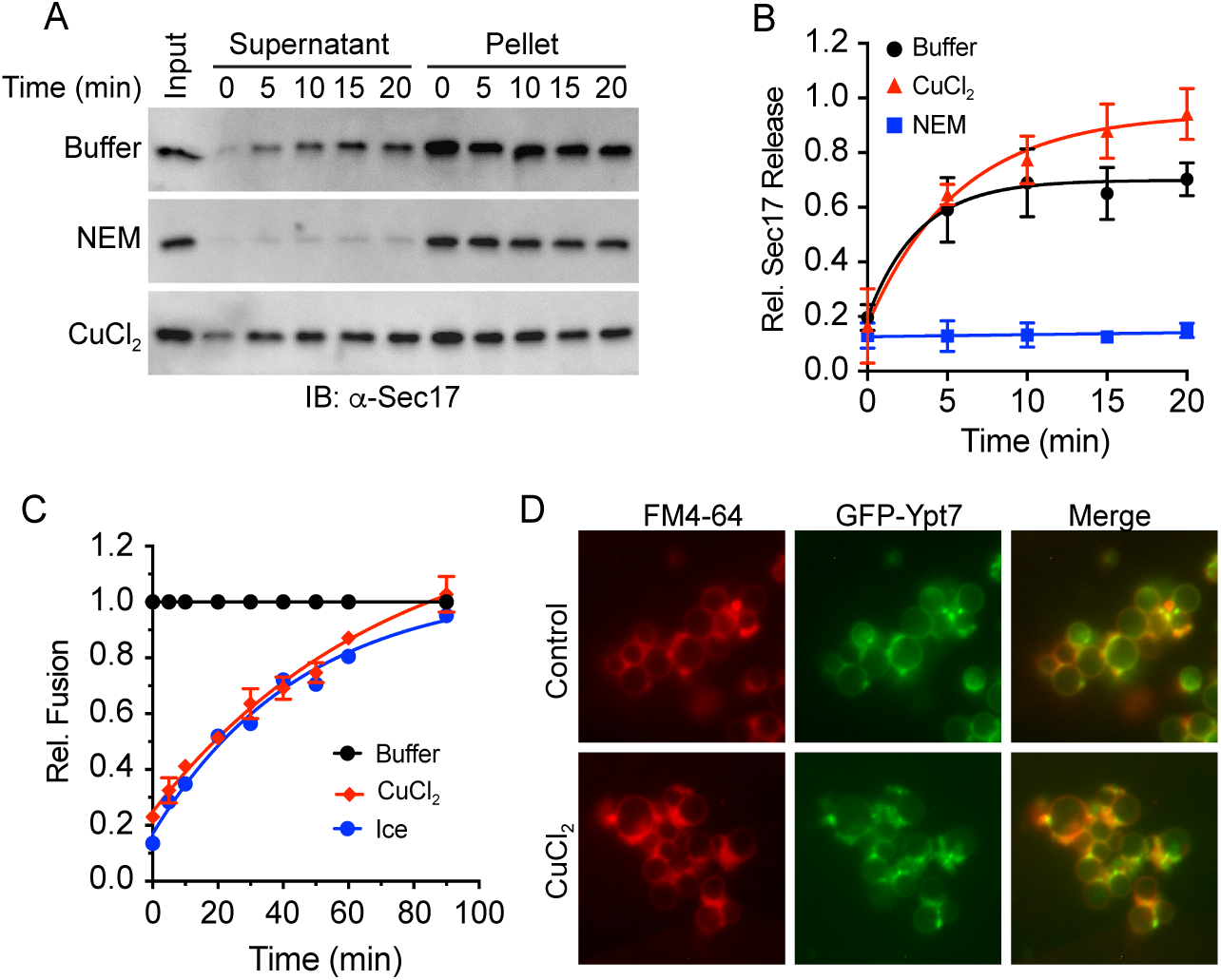
Cu^2+^ inhibits vacuole fusion after the docking stage. **(A)** Vacuoles from BJ3505 were monitored for the release of Sec17 from the membrane upon SNARE priming. Fusion reactions containing 3 µg of vacuoles (by protein) were incubated with reaction buffer, 100 µM Cu^2+^ or 1 mM NEM. Vacuoles were incubated at 27°C for the indicated times after witch the organelles were pelleted by centrifugation and solubilized proteins in the supernatant were separated from the membrane bound fraction. The membrane pellets were resuspended in volumes of reaction buffer equal to the supernatant. Both fractions were mixed with SDS loading buffer and resolved by SDS-PAGE. Sec17 was detected by immunoblotting and the amount released was calculated by densitometry. **(B)** Normalized values were averaged and plotted over time of incubation. **(C)** Gain of resistance kinetic vacuole fusion assays were performed in the presence of reaction buffer or 100 µM Cu^2+^. Reactions were incubated at 27°C or on ice for 90 min. Reagents were added at the indicated time points. Fusion inhibition was normalized to the reactions receiving buffer alone. Data were fit with first order exponential decay. Error bars are S.E.M. (n=3). **(D)** Isolated vacuoles harboring GFP-Ypt7 were incubated with or without 100 µM Cu^2+^ for 30 min at 27°C after which reaction tubes were placed on ice and labeled with FM4-64. Vacuoles were visualized by fluorescence microscopy.

To further resolve the stage of the fusion pathway that was inhibited by Cu^2+^ we performed a temporal gain of resistance assay (20–22). Individual fusion reactions were treated with inhibitors at different time-points throughout the incubation period. Vacuole fusion gains resistance to an inhibitor once the target of the reagent has had completed its function. For example, an inhibitor of SNARE priming will no longer be effective to anti-Sec17 antibody as the reaction enters a later stage such as hemifusion. With this test, Cu^2+^ was shown to continue inhibiting the fusion reaction after docking as its resistance curve lied on the ice curve **(Fig. 3C)**. Because this test only measures the last time an inhibitor functions, it was possible that Cu^2+^ could block stages between priming and full content mixing. This was in accord with the lack of an effect on vesicle docking and localization of Ypt7 at vertex sites **(Fig. 3D)**.

### Cu^2+^ inhibits trans-SNARE complex formation

Next, we examined if SNARE complex formation was affected by Cu^2+^. For this, we used a SNARE complex formation assay using exogenous GST-Vam7 in reactions treated with anti-Sec17 IgG to block priming (23–26). Thus, only the formation of new SNARE complexes will be measured in the presence of Cu^2+^. After the addition of anti-Sec17 we further treated individual reactions with GDI to block tethering or Cu^2+^. Next, GST-Vam7 was added, and the reactions were incubated for 70 min at 27°C or on ice as indicated. After incubation, the reactions were placed on ice and processed for SNARE complex isolation as described in the Experimental Procedures. GST-Vam7 SNARE complexes were bound to reduced glutathione beads and proteins were resolved by SDS-PAGE. Specific proteins were detected by Western blotting. As previously seen, GST-Vam7 formed complexes with its cognate SNAREs Vam3 and Nyv1 when incubated at 27°C, but not when incubation on ice prevented complex formation as seen previously (23, 24) **(Fig. 4A-B)**. As a negative control GDI inhibited SNARE complex formation. When 15 µM CuCl_2_ was added we observed nearly a 50% reduction in SNARE pairing. We were limited to using 15 µM CuCl_2_ due to its interference with the glutathione-GST interactions at higher concentrations. This is in accord with work showing that copper can inhibit GST function in the cytoplasm (27). Nevertheless, 15 µM CuCl_2_, which blocks fusion by 50% was still able to substantially inhibit SNARE complex formation.

**Figure 4.**
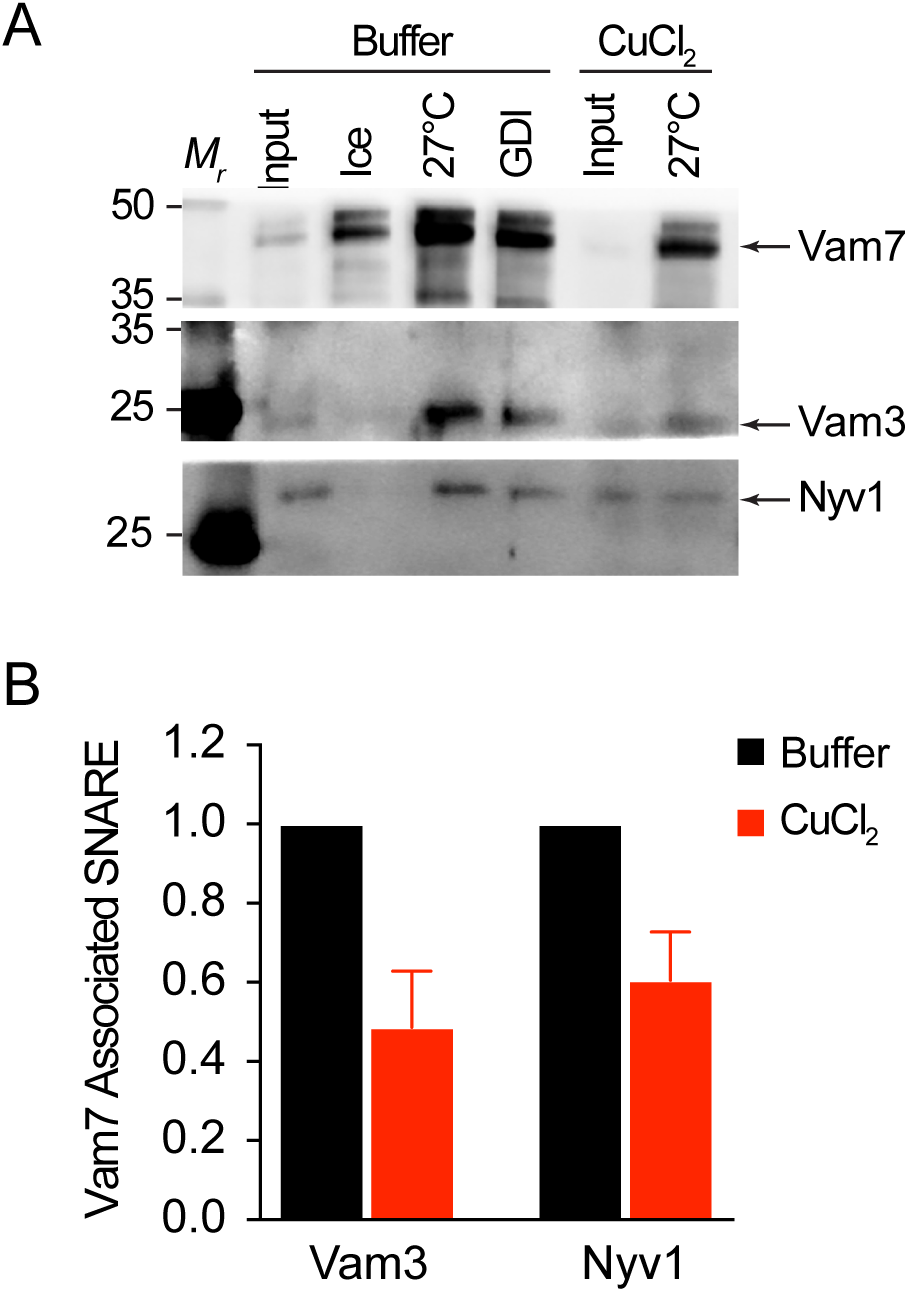
Cu^2+^ inhibits SNARE complex formation. **(A)** Large scale vacuole fusion reactions (6x) were incubated with anti-Sec17 IgG to block SNARE priming. After incubating for 15 min at 27°C, select reactions were further treated with either reaction buffer, 2 µM GDI or 15 µM Cu^2+^ and incubated for 5 min before adding 150 nM GST-Vam7. Reactions were then incubated for an additional 70 min. One reaction remained on ice for the duration of the assay. Reactions were then processed for glutathione pulldown of GST-Vam7 protein complexes. Isolated protein complexes were resolved by SDS-PAGE and probed for the SNAREs Vam3 and Nyv1. **(B)** Quantitation of SNARE complex formation in the presence or absence of CuCl_2_. Error bars are S.E.M. (n=3).

### Copper inhibits V-ATPase activity

Copper ions have been shown to inhibit the H^+^-ATPase activity in aquatic fungi (28), chromaffin cells (29), and the yeast plasma membrane H^+^-ATPase Pma1 (30). Because vacuole fusion is inhibited when the proton gradient is collapsed, indicating a role for V-ATPase function (31), we proposed that Cu^2+^ could inhibit vacuole fusion by blocking V-ATPase activity. To test this, we used an acridine orange (AO) fluorescence quenching assay to measure V-ATPase activity (32). Transport of protons into the vacuole lumen leads to the quenching of AO fluorescence. Fluorescence quenching was reversed when the proton gradient was disrupted with FCCP. Using this method, we found that ≥50 µM CuCl_2_ completely inhibited AO quenching indicating that V-ATPase activity was blocked **(Fig. 5A-B)**. To test whether other divalent cations also block V-ATPase activity we next tested the effects of Mg^2+^, Zn^2+^ and Co^2+^on the AO assay. In **Figure 5C** we show that none of the cations tested blocked proton transport even at 100 µM, suggesting that the effect of Cu^2+^ was specific. To further verify the effects of Cu^2+^ on V-ATPase activity we monitored quinacrine fluorescence of yeast vacuoles. Quinacrine accumulates and fluoresces in acidic compartments (33). Quinacrine staining showed that wild type cells stained with quinacrine while those treated with CuCl_2_ failed to accumulate quinacrine **(Fig. 5D)**. Thus, we have concluded that Cu^2+^ blocks vacuolar V-ATPase function.

**Figure 5.**
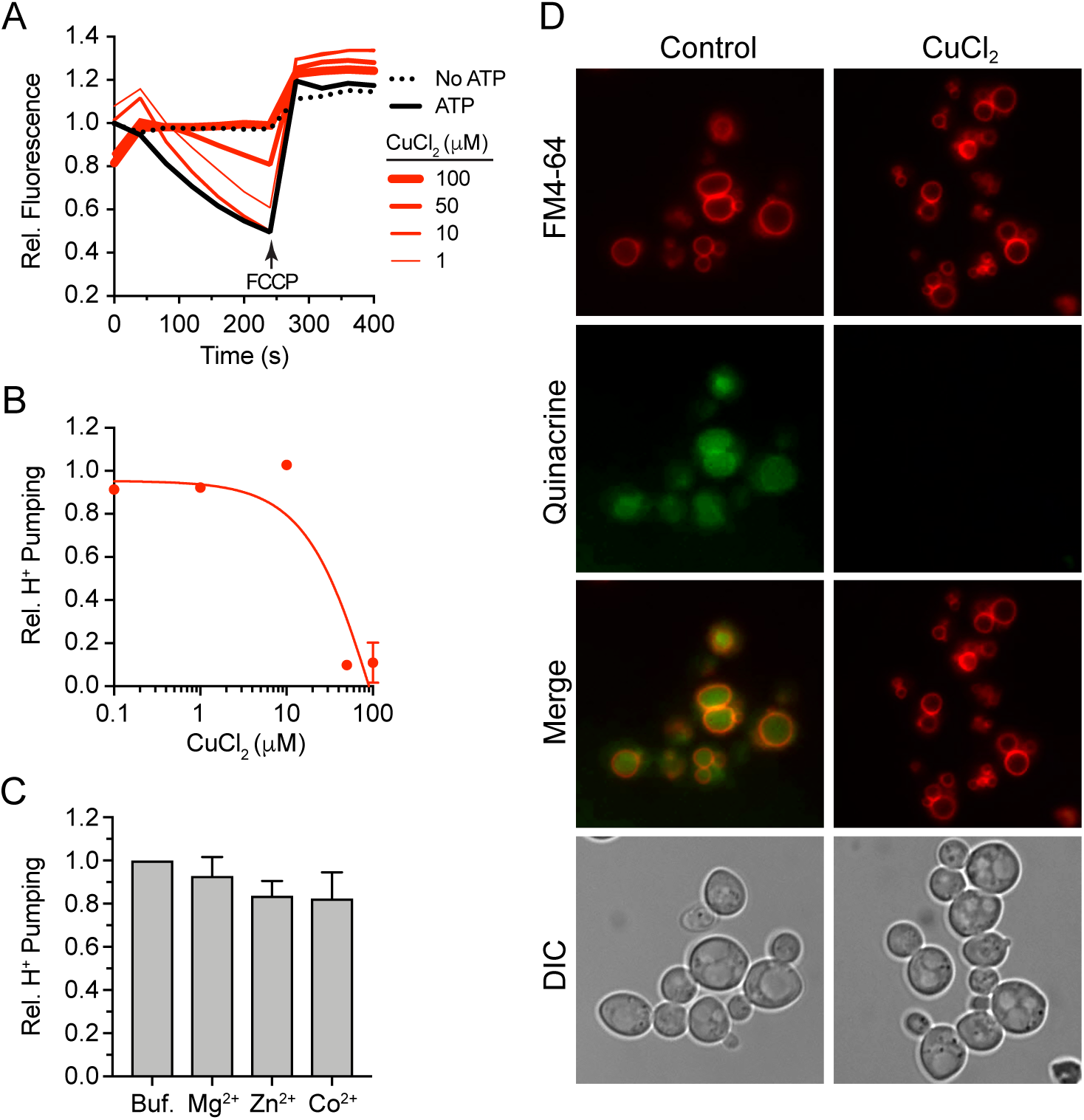
V-ATPase activity is blocked by Cu^2+^. **(A)** Acridine orange (AO) fluorescence quenching assays were used to monitor H^+^ pumping into the vacuole lumen. Vacuoles were incubated with dose curve of CuCl_2_ or buffer (± ATP) and 15 µM AO. AO fluorescence quenching was measured using a plate reader. Fluorescence was measured every 40 seconds and plotted against time. **(B)** Quantitation of average maximum AO fluorescence. **(C)** AO fluorescence quenching in the presence of 100 µM MgCl_2_, ZnCl_2_ or CoCl_2_. Error bars are S.E.M. (n=3). **(D)** Log phase BJ3505 cells were incubated with 50 µM CuCl_2_ for 1h. Vacuoles were stained with 2 µM FM4-64 and 200 µM quinacrine.

Taken together, this study has demonstrated that elevated Cu^2+^ concentrations are deleterious to the fusion machinery and the V-ATPase. Metals such as Cu^2+^ are often associated with generating ROS that can lead to membrane damage through lipid peroxidation. Damaging the vacuolar membrane would lead to the loss of a membrane potential and ion gradients that are essential for vacuole homeostasis. That said, the inhibitory effect of Cu^2+^ on vacuole fusion was independent of ROS and membrane damage. Instead we observed an effect on vacuole acidification and SNARE complex formation. The effects of Cu^2+^ on the VATPase and SNAREs are likely due to separate direct mechanisms. The disparity can be seen in the difference in the kinetics of resistance. In this study we showed that Cu^2+^ was a late inhibitor with a resistance curve that overlaid the ice curve. In comparison, the inhibition of SNARE pairing has been shown many times to occur much earlier during the docking stage the pathway with a halftime of 20 min (20, 34, 35).

While kinetically separate, the effects of Cu^2+^ on these functions could have a common origin such as the reduction of membrane fluidity. The mean by which Cu^2+^ and not Zn^2+^ or other metals (except for Ag^2+^) alter membrane fluidity is due copper-copper interactions when bound to anionic phospholipids (36). The interaction with anionic phospholipids reduces Cu^2+^ to Cu^+^ which stabilizes the complex. This occurs independent of a reductase. The formation of the copper-lipid complexes precludes chelation and is likely to be the cause for reducing membrane fluidity. Copper has been shown to reduce membrane fluidity in various cell types, organisms and artificial supported bilayers (7, 36–41). Thus, Cu ions could indirectly alter the vacuolar V-ATPase, as other H^+^/V^+^-ATPases have been shown to be inhibited when membrane fluidity is reduced (39, 41–44). Separately, we and others have found that SNARE function can be altered by membrane fluidity (24, 25, 45). Together, these notions and the data presented add another facet to pleiotropic effects of this metal to include membrane fusion.

### Experimental Procedures

#### Reagents

Soluble reagents were dissolved in PIPES-Sorbitol (PS) buffer (20 mM 1,4-piperazinediethane sulfonic acid (PIPES)-KOH pH 6.8 and 200 mM sorbitol) with 125 mM KCl unless indicated otherwise. Anti-Sec17 IgG (18), and Pbi2 (Protease B inhibitor) (46) were prepared as described previously. Recombinant GST-Vam7 and GDI were prepared as previously described (23, 24, 47) NEM (N-ethylmaleimide), acridine orange and FCCP were from Sigma. Quinacrine was from Cayman Biochemical.

#### Strains

BJ3505, DKY6281, RFY74 (BJ3505 *yvc1*Δ) and RFY75 (DKY6281 *yvc1*Δ) were previously described (21, 48, 49) were used for fusion assays. *FRE6* was deleted by homologous recombination using PCR products amplified from pAG32 with primers 5’-FRE6-KO (5’-ATCT TCTAAAGTGAAGCATGACGACCATAGCTCGTTGAATCTGTTTAGCTTGCCTTGTCC–3’) and 3’-FRE6-KO (5’-TATAGGTGGGCGTAGGATCAGAAGGAGCCGGAGAGAAGATGAC ACTGGATGGCGGCGTTA–3’). The PCR product was transformed into BJ3505 and DKY6281 yeast by standard lithium acetate methods and plated on YPD media containing hygromycin (200 µg/μl) to generate DKY6281 *fre6*Δ::*hphMX4* (RFY93) and BJ3505 *fre6*Δ*:: hphMX4* (RFY94).

#### Vacuole Isolation and in vitro fusion

Vacuoles were isolated as described (48). *In vitro* fusion reactions (30 µl) contained 3 µg each of vacuoles from BJ3505 (*PHO8 pep4*Δ) and DKY6281 (*pho8*Δ *PEP4*) backgrounds, reaction buffer 20 mM PIPES-KOH pH 6.8, 200 mM sorbitol, 125 mM KCl, 5 mM MgCl_2_), ATP regenerating system (1 mM ATP, 0.1 mg/ml creatine kinase, 29 mM creatine phosphate), 10 µM CoA, and 283 nM Pbi2. Fusion was determined by the processing of pro-Pho8 (alkaline phosphatase) from BJ3505 by the Pep4 protease from DK6281. Fusion reactions were incubated at 27°C for 90 min and Pho8 activity was measured in 250 mM Tris-HCl pH 8.5, 0.4% Triton X-100, 10 mM MgCl_2_, and 1 mM *p*-nitrophenyl phosphate. Pho8 activity was inhibited after 5 min by addition of 1 M glycine pH 11 and fusion units were measured by determining the *p-*nitrophenolate levels by measureing absorbance at 400 nm.

#### GST-Vam7 SNARE complex isolation

SNARE complex isolation was performed as described previously using GST-Vam7 (23, 24, 26, 50) Briefly, 5X fusion reactions were incubated with 85 µg/ml anti-Sec17 IgG to block priming. After 15 min, 2 µM GDI or 15 µM CuCl_2_ was added to selected reactions and incubated for an additional 5 min before adding 150 nM GST-Vam7. After a total of 90 min, reactions were sedimented (11,000 *g*, 10 min, 4°C), and the supernatants were discarded before extracting vacuoles with solubilization buffer (SB: 20 mM HEPESKOH, pH 7.4, 100 mM NaCl, 2 mM EDTA, 20% glycerol, 0.5% Triton X-100, 1 mM DTT) with protease inhibitors (1 mM PMSF, 10 µM Pefabloc-SC, 5 µM pepstatin A, and 1 µM leupeptin). Vacuole pellets were overlaid with 100 µl SB and resuspended gently. An additional 100 µl SB was added, gently mixed, and incubated on ice for 20 min. Insoluble debris was sedimented (16,000 *g*, 10 min, 4°C) and 176 µl of supernatants were removed and placed in chilled tubes. Next, 16 µl was removed from each reaction as 10% total samples, mixed with 8 µl of 3X SDS loading buffer and heated (95°C, 5 min). Equilibrated glutathione beads (30 µl) were incubated with the remaining extracts (15 h, 4°C, nutation). Beads were sedimented and washed 5X with 1 ml SB (735 *g*, 2 min, 4°C), and bound material was eluted with 40 µl 1X SDS loading buffer. Protein complexes were examined by Western blotting.

#### Proton Pumping

The proton pumping activity of isolated vacuoles was performed as described by others with some modifications (32). *In vitro* H^+^ transport reactions (60 µl) contained 20 µg vacuoles from BJ3505 backgrounds, fusion reaction buffer, 10 µM CoA, 283 nM Pbi2, and 15 µM of the H^+^ probe acridine orange. Reaction mixtures were loaded into a black, half-volume 96-well flat-bottom plate with nonbinding surface. ATP regenerating system or buffer was added and reactions were incubated at 27°C while acridine orange fluorescence was monitored. Samples were analyzed in a fluorescence plate reader with the excitation filter at 485 nm and emission filter at 520 nm. Reactions were initiated with the addition of ATP regenerating system following the initial measurement. After fluorescence quenching plateaus we added 30 µM FCCP to collapse the proton gradient and restore acridine orange fluorescence.

#### Vacuole docking

Docking reactions (30 µl) contained 6 µg of vacuoles from GFP-Ypt7 yeast were incubated in docking buffer (20 mM PIPES-KOH pH 6.8, 200 mM sorbitol, 100 mM KCl, 0.5 mM MgCl_2_), ATP regenerating system (0.3 mM ATP, 0.7 mg/ml creatine kinase, 6 mM creatine phosphate), 20 μM CoA, and 283 nM Pbi2 (Protease B inhibitor) (51). Reactions were treated with 100 µM CuCl_2_ or buffer alone for 1 h at 27°C. After incubating, reaction tubes were placed on ice and vacuoles were stained with 1 µM MDY-64 and mixed with 50 μl of 0.6% low-melt agarose. Samples were vortexed to disrupt non-specific clustering, mounted on slides and imaged by fluorescence microscopy. Images were acquired using a Zeiss Axio Observer Z1 inverted microscope equipped with an X-Cite 120XL light source, Plan Apochromat 63X oil objective (NA 1.4), and an AxioCam CCD camera.

## Author contributions

GEM, KDS and RAF conceived the project and designed experiments. GEM, KDS, AG, CZ, BCJ, LRH, MLS and DAR-K performed the experiments and analyzed data. RAF supervised the research. GEM, KDS and RAF wrote the manuscript with input from all authors.

## Acknowledgements

This research was supported by grants from the National Science Foundation (MCB 1818310 to RAF).

## Competing interests

The authors declare that they do not have conflicts of interest with the contents of this article.

## Footnotes

The abbreviations used:

AO: acridine orange;
BCS: Bathocuproine disulfonate;
FCCP: 2-[2-[4-(trifluoromethoxy) phenyl]hydrazinylidene]-propanedinitrile;
GDI: guanidine dissociation inhibitor;
NEM: N-ethylmaleimide;
PBN: *N-tert*-butyl-a-phenylnitrone;
SNARE: soluble N-ethylmaleimidesensitive factor attachment protein receptor;
TETA: Triethylenetetramine.

